# A novel IKK- and proteasome-independent mechanism of RelA activation triggers senescence associated secretome via transcriptional repression of NFKBIA

**DOI:** 10.1101/2019.12.19.882225

**Authors:** Marina Kolesnichenko, Nadine Mikuda, Uta E. Höpken, Maja Milanovic, A. Bugra Tufan, Bora Uyar, Wei Sun, Kolja Schleich, Linda von Hoff, Michael Willenbrock, Inge Krahn, Sabine Jungmann, Michael Hinz, Altuna Akalin, Soyoung Lee, Ruth Schmidt-Ullrich, Clemens A. Schmitt, Claus Scheidereit

## Abstract

The IκB kinase (IKK) - NF-κB pathway is activated as part of the DNA damage response and controls both resistance to apoptosis and inflammation. How these different functions are achieved remained unknown. We demonstrate here that DNA double strand breaks elicit two subsequent phases of NF-κB activation *in vivo* and *in vitro*, which are mechanistically and functionally distinct. RNA-sequencing reveals that the first phase controls anti-apoptotic gene expression, while the second drives expression of senescence-associated secretory phenotype (SASP) genes. The first, rapidly activated phase is driven by the ATM-PARP1-TRAF6-IKK cascade, which triggers proteasomal destruction of IκBα and is terminated through IκBα (*NFKBIA*) re-expression. The second phase is activated days later in senescent cells but is independent of IKK and the proteasome. An altered phosphorylation status of p65, in part driven by GSK3β, results in transcriptional silencing of *NFKBIA* and IKK-independent, constitutive activation of NF-κB in senescence. Collectively, our study reveals a novel physiological mechanism of NF-κB activation with important implications for genotoxic cancer treatment.

## INTRODUCTION

Chemo- and radiotherapies, activated oncogenes and shortened telomeres trigger via the DNA damage response (DDR) a terminal proliferative arrest called cellular senescence (Blagosklonny, 2014; Lasry & Ben-Neriah, 2015; Lee & Schmitt, 2019; Salama et al, 2014). The associated alterations include formation of senescence associated heterochromatin foci (SAHF), increased expression of cell-cycle inhibitors, including p21 (*CDKN1A*) and p16 (*CDKN2A*) and of inflammatory cytokines and chemokines that constitute the senescence associated secretory phenotype (SASP) and a related, low-grade inflammation termed senescence inflammatory response (SIR) that affects surrounding tissues in a paracrine manner (Lasry & Ben-Neriah, 2015; Shelton et al, 1999). The epigenetically controlled cell-cycle cessation serves as a cell-autonomous barrier to tumor formation (Braig et al, 2005; Collado et al, 2005; Reimann et al, 2010). Therefore, induction of senescence was considered as important in treating cancer and other pathologies. The inflammatory response, however, comprises factors that may instigate oncogenic transformation, cell migration, and cancer stemness (Acosta et al, 2013; Acosta et al, 2008; Chien et al, 2011; Freund et al, 2011; Hoare et al, 2016; Jing et al, 2011; Milanovic et al, 2018; Reimann et al, 2010; Salama et al, 2014).

The majority of SASP factors are transcriptional targets of NF-κB (Acosta et al, 2008; Chien et al, 2011; Freund et al, 2011; Jing et al, 2011; Kuilman et al, 2008; Lasry & Ben-Neriah, 2015). Although it is well established that NF-κB drives inflammatory gene expression in senescence, whether it also contributes to cell-cycle arrest remained unclear. DNA double strand breaks lead to rapid activation of NF-κB RelA/p65-p50, the most prevalent heterodimer (Smale, 2012). A signaling cascade that is activated by *ataxia telangiectasia mutated* (ATM) and poly(ADP-Ribose) polymerase 1 (PARP1), and also depends on TNFR-associated factor 6 (TRAF6), converges on the IκB kinase (IKK) complex (Hinz et al, 2010; Stilmann et al, 2009; Wu et al, 2006). The latter is composed of the regulatory IKKγ subunit and the kinases IKKα and IKKβ (Hayden & Ghosh, 2012; Hinz & Scheidereit, 2014), which phosphorylate IκBα, targeting it for degradation by the 26S proteasome. Liberated NF-κB translocates to the nucleus and activates transcription of its target genes, including *NFKBIA,* encoding IκBα (Hinz et al, 2012). This negative feedback loop ensures that NF-κB activation is transient. Replication-, oncogene- and therapy-induced senescence is associated with unresolved DNA damage and with constitutive NF-κB activation (Chien et al, 2011; Freund et al, 2011; Jing et al, 2011; Rodier et al, 2009).

Using cells of epithelial origin, both from transgenic mouse models and from human primary and cancer cell lines, we demonstrate here that DNA damage triggers two functionally distinct phases of NF-κB activation. Whereas the first, immediately activated phase is IKK- and proteasome-dependent and activates anti-apoptotic gene expression, the second NF-κB activation phase, occurring days later in senescence, is caused by a permanent silencing of *NFKBIA* transcription and is thus IKK- and proteasome-independent. We show that this second, IKK-independent phase of NF-κB is responsible for SASP, but not for the cell cycle arrest. Furthermore, SASP expression profile is generated *in vivo* and *ex vivo* in the absence of DNA damage solely by depletion of IκBα.

## RESULTS

### DNA damage activates NF-κB d in two distinct phases: A transient anti-apoptotic first and a persistent inflammatory second phase

To investigate the kinetics of activation of NF-κB, we first examined the establishment of senescence and SASP over time in human diploid fibroblasts (HDFs) and cancer cell lines that experienced DNA damage. Onset of senescence was marked by senescence-associated β-galactosidase (SA-β-gal) activity and elevated p21CIP1 expression (Fig 1A and Appendix Fig S1A). Proliferation ceased 1-2 days following irradiation (IR), as expected, and cells entered a lasting senescent state (Appendix Fig S1B and data not shown). Unresolved DNA damage, evidenced by γH2AX foci, peaked within minutes and persisted through all time points (Appendix Fig S1C). Strikingly, the single dose IR generated a biphasic NF-κB activation, with two temporally separate phases of nuclear translocation and DNA binding, first, within hours and then days later (Fig 1B, and Appendix Fig S1C-E). An RNA-seq analysis revealed distinct transcriptomes in both NF-κB phases (Fig 1C and Tables EV1A-C). During the first phase, 289 transcripts showed significant upregulation (Table EV1A), which included direct targets of NF-κB, such as early response genes and negative feedback inhibitors of the pathway (NFKBIA and TNFAIP3; see Table EV1B). This transcript group was enriched for GO terms “cell cycle arrest” and “regulation of apoptotic process” (Table EV2A). In contrast, the 2,979 transcripts upregulated during the second phase were enriched for GO terms “inflammatory response”, “immune response”, and “response to wounding” (Fig 1C and Table EV2B). The biphasic NF-κB response thus coincided with two distinct transcriptomes, an anti-apoptotic first phase and a pro-inflammatory second phase.

**1.**
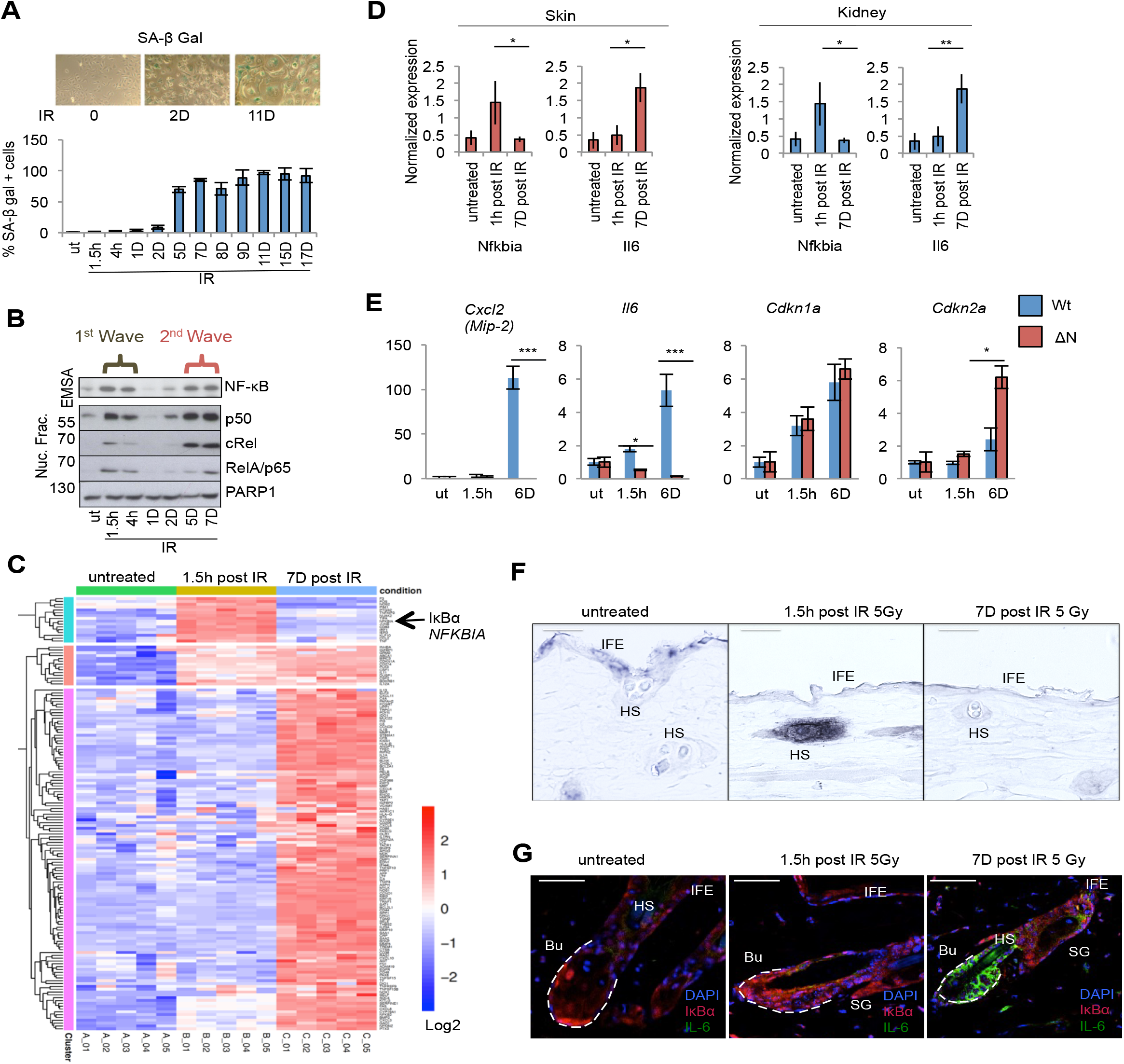
NF-κB is activated in two distinct phases in response to DNA damage corresponding to anti-apoptotic and pro-inflammatory gene programs. (A) SA-β-gal staining of U2-OS cells, harvested at indicated times post irradiation (IR); h, hours; D, days; UT, untreated. N=3. (B) Nuclear fractions from U2-OS cells were harvested at indicated time points post IR (10 Gy) as in (A). Top panel: NF-κB DNA binding was analyzed by EMSA. Lower panels: Western blot of indicated NF-κB subunits and PARP1 as loading control. The two phases of nuclear translocation and DNA binding are indicated. (C) Gene expression analysis of two NF-κB phases following damage. RNA-Seq analysis (quintuplicate samples) of U2-OS cells, either untreated or analyzed 1.5 hours or 7 days after irradiation (10 Gy), as indicated. The heatmap shows significantly regulated genes (log2 value > 0.5 and p value < 0.05). (D) Mice (triplicate per condition) were irradiated (5 Gy) and sacrificed after 1 hour (first phase), 7 days (second phase) or left untreated. Skin or renal tissue RNA was analyzed by qRT-PCR, shown as a mean +/− SD. Significance was confirmed by unpaired T test * < p 0.05 for skin and SD * = p < 0.05, ** = p < 0.01 for kidney samples, respectively. (E) Kidney cells from IκBα^ΔN^ mice (ΔN, N=2) or control littermates (Wt, N=2) were irradiated either 1.5h or six days prior to harvest or left untreated (ut). RNA was analyzed by qRT-PCR for indicated genes. (F) *In situ* hybridization using an IκBα mRNA probe on skin sections from mice treated as in (D) for the time indicated. For abbreviations, see (F). (G) Skin sections as in (D) analyzed by immunofluorescence with IκBα (red) or IL-6 antibody (green) and nuclear DAPI staining (blue). Dotted lines delineate hair follicles, Bu, bulge region, HS, hair shaft, IFE, interfollicular epidermis, SG, sebaceous gland.

We next analyzed murine tissues for IR-induced expression of the representative first- and second-phase target genes of NF-κB, *Nfkbia* and *Il6*, respectively. IR strongly activated NF-κB in kidney and skin (unpublished data). Indeed, IR significantly induced *Nfkbia* mRNA in both tissues only at early and *Il6* mRNA only at late time points, representing the first and second NF-κB phases (Fig 1D).

To ensure that the expression of SASP during the second phase depends on NF-κB, we analyzed primary kidney cells from irradiated mice, which ubiquitously express the NF-κB super-repressor IκBαΔN (Krappmann et al, 1996; Schmidt-Ullrich et al, 2001). Compared to littermate controls, expression of *Cxcl2* and *Il6* in the second phase was abolished, whereas *Cdkn1a* and *Cdkn2a* upregulation was unaffected (Fig 1E). These *in vivo* results further reveal that DNA damage activates NF-κB that drives SASP, but NF-κB is not essential for the proliferative arrest observed in senescence.

A strong response was seen in hair follicles (HF), which require NF-κB activation for development and morphogenesis (Schmidt-Ullrich et al, 2001). Upregulation of *Nfkbia* mRNA and IκBα protein was restricted to the first phase of NF-κB activation in HF following whole body IR (Fig 1F and 1G). At 7 days post IR, IκBα expression in the proximal HF was strongly reduced, concomitant with an increase in IL-6 expression in the same region (Fig 1G). These results demonstrate that two distinct, subsequent NF-κB transcriptomes also occur *in vivo*. Because IκBα is required to terminate NF-κB signaling, we postulated that loss of IκBα in senescence could trigger SASP.

### Loss of IκBα expression in senescence triggers the second phase of NF-κB activation and generates SASP

IκBα expression was either undetectable or strongly reduced in senescence in the different epithelial cancer- and non-transformed cell lines tested (Fig 2A and Appendix Fig S2A-D). Likewise, *NFKBIA* mRNA was upregulated only in the first phase (Fig 2A, right panel; Appendix Fig S2E), despite robust activation of NF-κB in both phases (Fig 2B, lanes 2 and 4). Repeated IR treatment of senescent cells restored neither *NFKBIA* mRNA nor IκBα protein expression (Fig 2A left, lane 5, and right panel).

**Figure 2.**
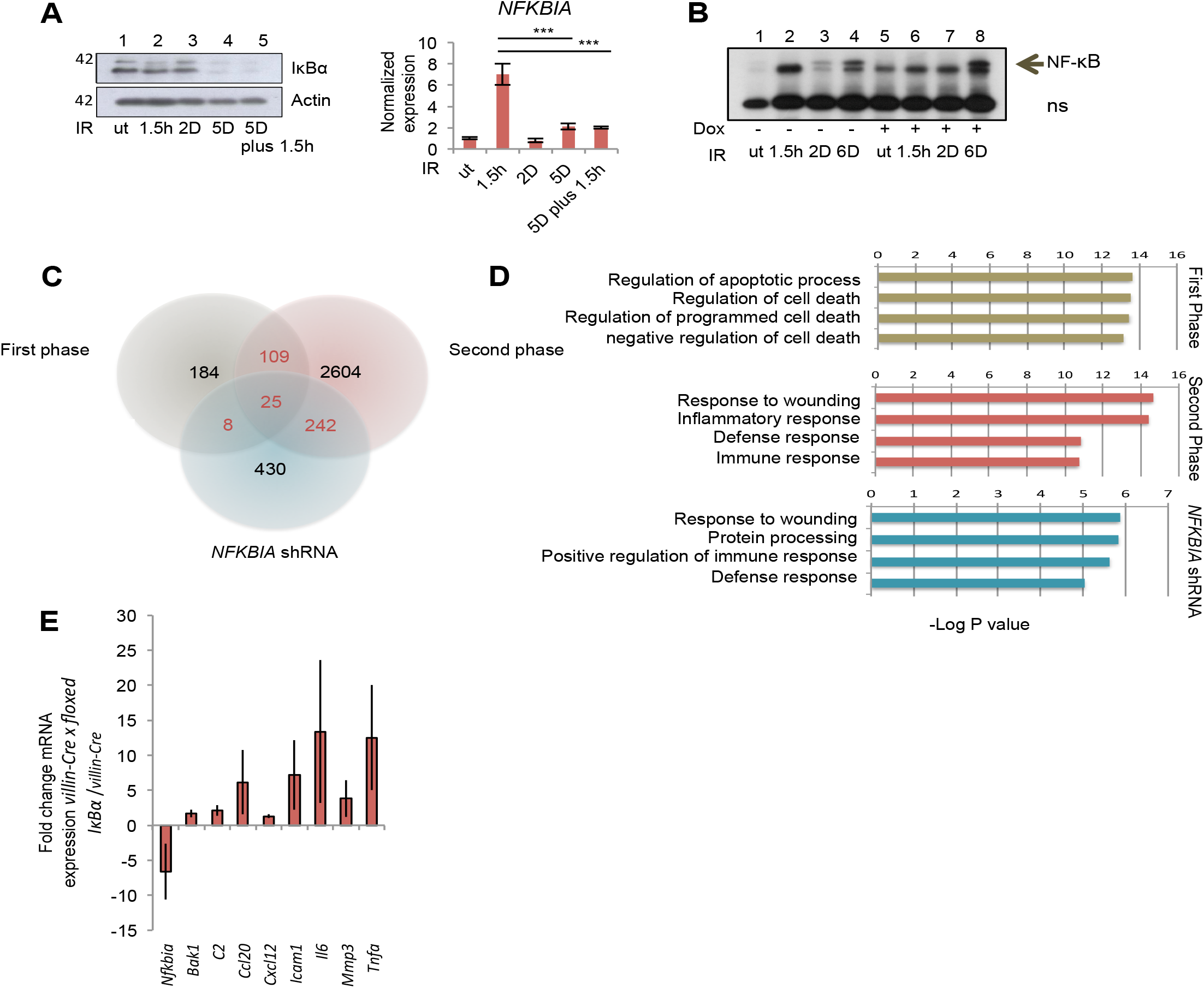
Knockdown of *NFKBIA* in epithelial cells and knockout *in vivo* mimics second phase NF-κB activity and triggers SASP. (A) Left panel: IκBα western blot of U2-OS cells. Lane 1, untreated (ut); Lanes 2-4, single IR (20 Gy) at indicated time points prior to harvest; Lane 5, IR at 5 days plus 1.5 hours prior to harvest. Right panel: Samples treated as above, analyzed by qRT-PCR. (B) *NFKBIA* knockdown by dox-inducible shRNA in U2-OS cells. Cells were irradiated or not, as indicated and described above. NF-κB activity was analyzed by EMSA. ns, non-specific band. (C) Venn diagram of RNA-Seq analysis of genes activated following irradiation in the first and second NF-κB activity phases or following shRNA-mediated *NFKBIA* knockdown in non-irradiated cells (as in B). (D) Top GO terms obtained using DAVID Functional Annotation Bioinformatics Microarray 6.7 for first and second phase genes and genes activated upon *NFKBIA* knockdown, as in (C), as indicated. (E) qRT-PCR analysis of indicated genes was performed using RNA extracted from the duodenum of 5 and 8 weeks old control *villin-Cre* (N=6) or *villin-Cre x floxed IκBα* mice (N=6). Expression is shown as fold change between littermates, paired T-test. * P< 0.05 ** = P < 0.01, and # P = 0.05.

Activated oncogenes cause DNA strand breaks and induce DDR signaling, thereby promoting cellular senescence similar to cells exposed to DNA-damaging agents (Acosta et al, 2008; Coppe et al, 2008; Kuilman et al, 2008). Inducible activation of oncogenic RASV12 led to activation of NF-κB and expression of the representative SASP factor IL-8 (encoded by *CXCL8*) that negatively correlated with IκBα expression (Appendix Fig S2F-G). In summary, these data show that different pro-senescent triggers lead to loss of *NFKBIA* mRNA expression together with the onset of an NF-κB-driven SASP.

To investigate if experimental *NFKBIA* depletion would mimic the second phase NF-κB activation and SASP type gene expression, we performed *NFKBIA* knockdown experiments (Fig 2B and Appendix Fig S2H-I). Proliferation of cells was unaffected by knockdown of NFKBIA (Appendix Fig S2J). However, untreated *NFKBIA*-depleted cells that had not experienced DNA damage, revealed increased NF-κB DNA-binding activity (Fig 2B, compare lanes 1 and 5), comparable to the levels observed in irradiated, senescent cells at day 6 (compare lanes 4 and 5). IκBα rescue through ectopic overexpression of proteasome-insensitive mutant S32AS36A blocked the induction of the bona-fide SASP factors (Fig S2J), consistent with the conclusion that loss of IκBα drives expression of SASP.

Remarkably, the transcriptome of non-irradiated *NFKBIA*-depleted cells shared a strong overlap and the same GO terms with that of the second phase, senescent cells (Fig 2C and D, and Table EV2B-C). Thus, loss of IκBα in the absence of DNA damage is sufficient to generate SASP. Secretion of a set of the identified cytokines and chemokines was confirmed by an antibody array (Appendix Fig S3A). Some of these, including IL-6 and GM-CSF, activate monocytes and migration of macrophages (Bachelerie et al, 2014). In fact, supernatants from either senescent or IκBα-depleted cells induced migration of macrophages (Appendix Fig S3B).

We next asked whether knockout of *Nfkbia* would also trigger SASP *in vivo*. Since ubiquitous loss of IκBα expression causes early postnatal lethality (Beg et al, 1995; Klement et al, 1996), we generated intestinal epithelium-restricted (*villin-Cre* x floxed *Nfkbia*) knockout mice. The expression of selected typical SASP factors, including known NF-κB targets *Ccl20*, *ICcam1*, *Il6* and *Tnfa*, was indeed activated in the small intestines of knockout compared to control *villin-Cre* littermates (Fig 2E), confirming our conclusion in an *in vivo* setting.

### IκBα loss in senescence is mediated by posttranslational modifications of p65

Since we observed distinct NF-κB targets as upregulated during the two phases (Table EV1B), we next knocked down p65 to determine whether the same family member is responsible for both transcriptomes. All NF-κB targets, including SASP, showed decrease in expression, in cells bearing shRNA against *RELA*/p65 (Fig 3A). Nevertheless, knockdown of *RELA* did not rescue cells from senescence (Fig 3B), indicating that it does not contribute to cell cycle arrest.

**Figure 3.**
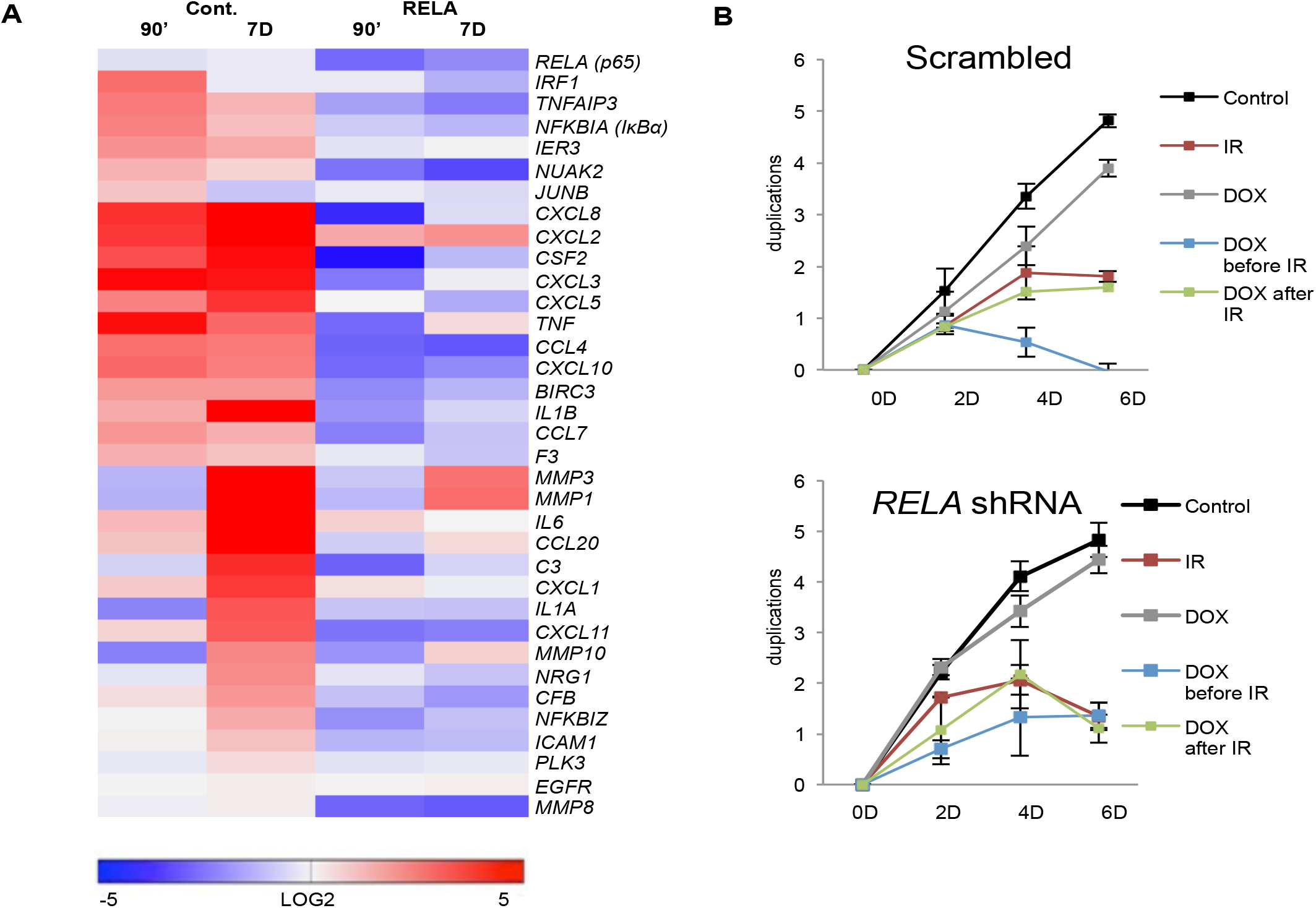
RelA regulates SASP but not cell cycle arrest. (A) Expression of SASP targets of NF-κB obtained from RNA-seq analysis (Supplemental Table 1B), was quantitated using qRT-PCR from the shRNA (*RELA*) stably expressing U2-OS cells. Cells were treated with doxycycline (Dox) to induce knockdown of p65. Heatmap represents targets normalized to the untreated Scrambled control. Expression is shown as log2 change with p values < 0.05. First-wave and second-wave samples were irradiated 90 minutes and seven days prior to harvest, respectively. (B) Cell duplication was measured at time points indicated in U2-OS cells bearing either scrambled control or Dox-inducible shRNA against *RELA* (biological triplicates). Treatment with Dox was initiated at two days prior to IR (10 Gy) or after IR.

Since loss of IκBα in the second phase resulted from decline of mRNA expression, we investigated its regulation by p65. We found that p65 was recruited to the NFKBIA (IκBα) promoter in both first- and second-phases of activation with similar efficiencies (Appendix Fig S4A). We next investigated the phosphorylation status of p65 during the two phases (Fig 4A). Phosphorylation on p65 Ser536, the substrate site of IKK, peaked during the first phase and declined in the second. Unlike phosphorylation of p65 at S536 and S267, which enhance the p65 transactivation potential, phosphorylation at S468 is inhibitory and is mediated in part by GSK3β (Buss H, 2004; Christian F, 2016). Of note, GSK3β exhibits increased kinase activity in senescence to activate formation of SAHF, through downregulation of Wnt signaling (Ye et al, 2007). Indeed, nuclear phosphorylation on S468 increased in senescence (Fig 4A left and right panels). To determine if phosphorylation on S468 repressed expression of IκBα, we overexpressed p65 bearing a S468A mutation in cells where endogenous p65 was knocked down. Overexpression of p65 S468A rescued IκBα expression. Similar results were observed with ectopic expression of wildtype p65, likely due to the abundance of the substrate in relation to the S468 kinase(s). As a negative control, we transfected the S267A mutant. Since acetylation at S267 is required for p65 activity (Christian F, 2016), as expected, its overexpression did not rescue IκBα in senescence (Fig 4B).

**Figure 4.**
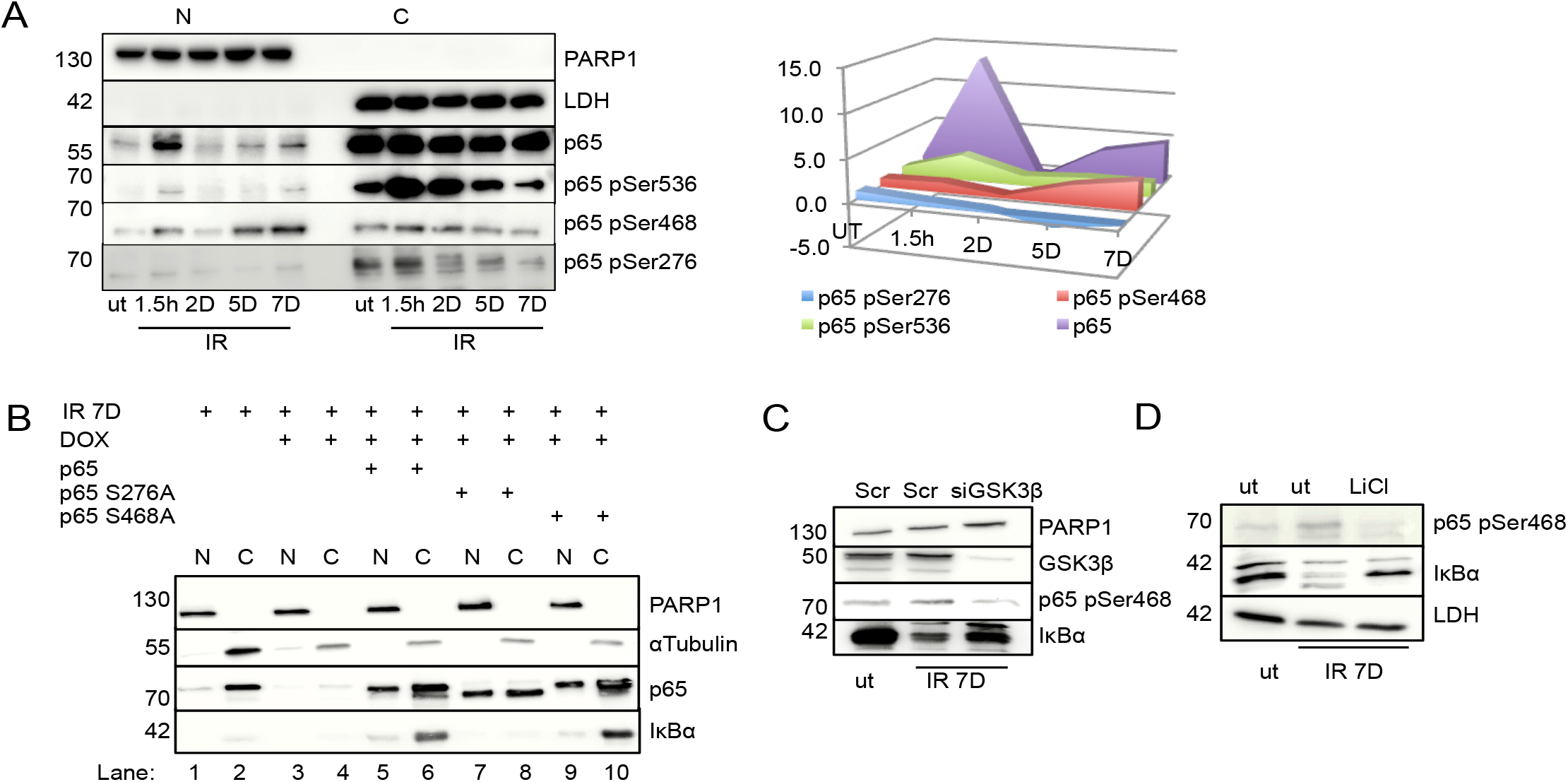
p65 Ser 468 phosphorylation in senescence inhibits *NFKBIA* expression. (A) Nuclear, N, and cytoplasmic, C, fractions of U2-OS cells were analyzed at the times after IR exposure, as indicated, by SDS-PAGE western blotting. PARP1 and LDH are fractionation controls and phospho-specific p65 signals are indicated. Right panel: quantitation of nuclear p65 and phosphorylated p65 species. Fold changes compared to untreated samples (ut) and time points post IR are indicated on Y and X axes, respectively. (B) U2-OS cells treated with dox to deplete endogenous p65, were irradiated (20 Gy) and transfected with plasmids encoding p65, p65-S276A or p65-S468A, as indicated. Nuclear (N) and cytoplasmic (C) lysates were analyzed by SDS-PAGE at day seven post IR.

To determine directly to which extent GSK3β contributes to phosphorylation of S468, we knocked down GSK3β or inhibited GSK3β by lithium chloride. Both modes of interference diminished S468 phosphorylation and led to a partial restoration of IκBα expression in senescence (Fig 4C and D). These data show that changes in p65 phosphorylation contribute to an attenuation of IκBα expression in senescence.

### Second phase NF-κB activation in senescence is independent of IKK-signaling and the proteasome

We next investigated the contribution of the known regulators of the genotoxic stress induced IKK pathway (Fig 5A) in the two NF-κB phases. The activation of ATM and IKK correlated only with the first NF-κB phase, with rapid IR-induction in the first hours followed by a decline afterwards (Fig 5B). We also found that TRAF6 depletion only abrogated activation of the first, but not the second NF-κB phase (Fig 5C). Expression of target genes of NF-κB, IL6 and IL1A, in the second phase were completely unaffected by depletion of PARP1 or ATM (Appendix Fig S4B and data not shown). Furthermore, the proteasome inhibitors Bortezomib and MG132 inhibited IR-induced IκBα destruction in the first NF-κB phase, but not loss of IκBα in the second phase (Fig 5B). In line with this, upregulation of the *bona-fide* SASP component IL-1α was not affected by proteasome inhibition (Fig 5D).

**Figure 5.**
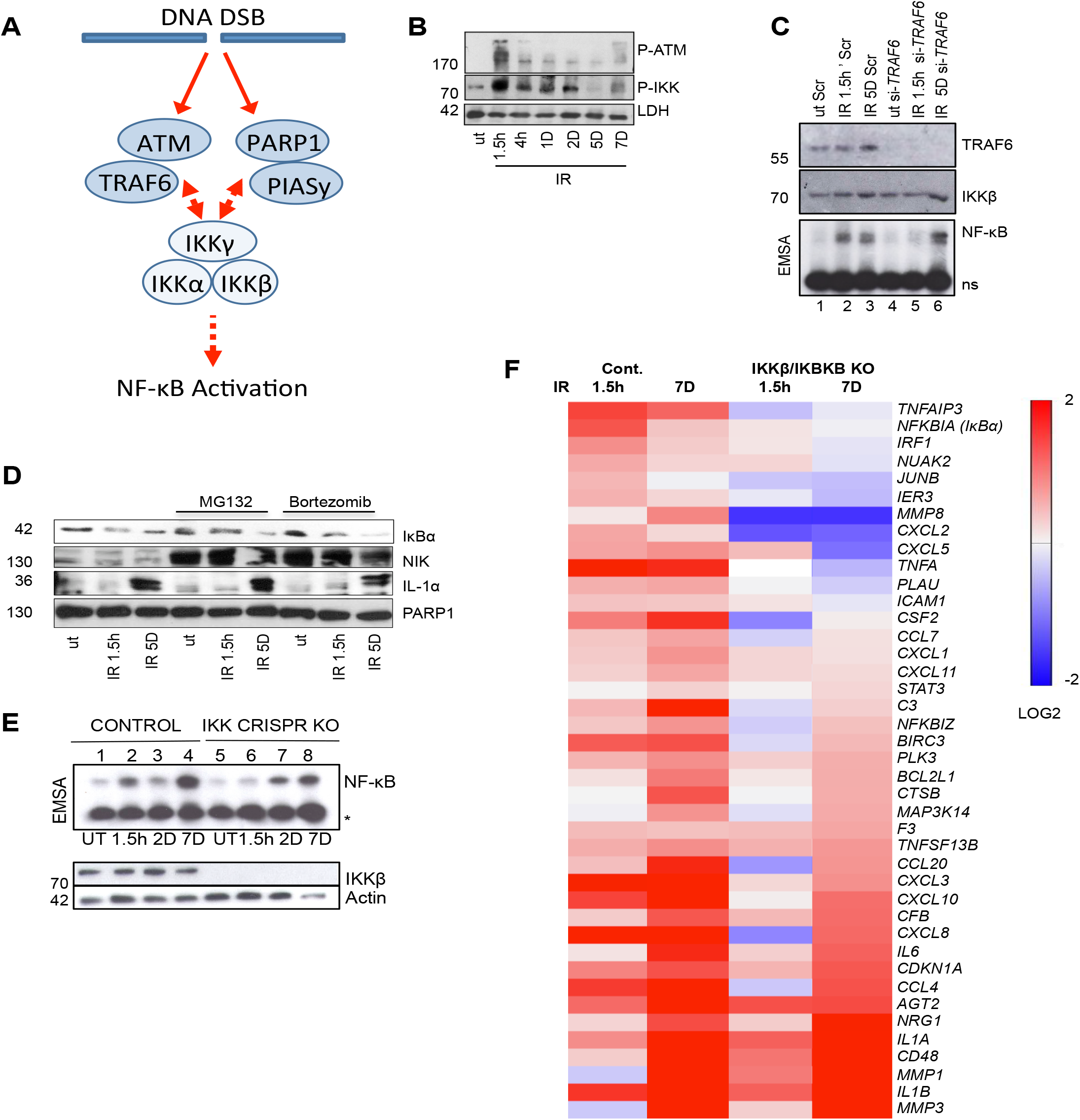
The second phase is IKK- and proteasome-independent. (A) Diagram showing DNA double strand break induced first phase NF-κB activation through an ATM/TRAF6 and PARP1/PIASy dependent IKK activation mechanism (Hinz et al, 2010; Stilmann et al, 2009; Wu et al, 2006). (B) SDS-PAGE western blot analysis of whole cell lysates of U2-OS cells irradiated at time points indicated prior to harvest. Molecular weight markers are indicated. ut, untreated. LDH, lactate dehydrogenase. (C) U2-OS cells transfected with *TRAF6* siRNA or scrambled control siRNA and TRAF6 and IKKβ levels analyzed by western blotting (top panels). Molecular weight marker positions are indicated. NF-κB activity was determined by EMSA in untreated cells (ut) and after IR at indicated time points (lower panel). ns, unspecific band. (D) U2-OS cells were exposed to IR at indicated time points prior to harvest and analyzed by western blotting for levels of IκBα, NIK, IL-1α and PARP1. Treatment with proteasomal inhibitors MG132 or bortezomib started 4 hours prior to harvest. NIK serves as positive control for efficient proteasome inhibition. Molecular weight marker positions are indicated. (E) U2-OS CRISPR I*KBKB* knockout and control cell lines were irradiated (20 Gy) and harvested at indicated time points. NF-κB activity was analyzed by EMSA (upper panel). Asterisk indicates an unspecific band. Lower panel, western blot analysis of actin and IKKβ. Molecular weight markers are indicated to the left. (F) Irradiated U2-O2 control and *IKBKB* knockout cells (as above) were analyzed by qRT-PCR at the time points indicated for expression of NF-κB target genes identified by RNA-Seq (Table EV1). (C) U2-OS cells were transfected with siRNA against GSK3β or scrambled control (Scr) and further untreated (ut) or irradiated 7 days before harvesting, as indicated. Whole cell lysates were analyzed by SDS-PAGE and western blotting. (D) U2-OS cells were left untreated or exposed overnight to 10 mM LiCl, with or without prior irradiation, as indicated, and analyzed as in (D).

We eliminated IKKβ expression using CRISPR/Cas9 (Fig 5E). Loss of IKKβ impeded only first phase NF-κB activation (Fig 5E, lanes 2 and 6). Remarkably, IKKβ was not required for the second phase of NF-κB activation (Fig 5E, lanes 4 and 8). Likewise, siRNA-mediated IKKγ depletion only diminished p65 nuclear translocation at 1.5 h, but not 5 days following IR (Appendix Fig S4C).

Our conclusion was further corroborated by identification of genes that depended on p65 expression (Fig 3), yet showed unaltered expression in senescent cells with CRISPR mediated knockout of *IKBKB* (Fig 5F). These IKK-independent genes included many bona fide SASP factors, such as IL-1α, IL-1β, IL-6, IL8 (Fig 5F, lower half of the panel). All these data substantiate that only the first phase of NF-κB depends on the IKK cascade induced by DNA damage and on proteasomal destruction of IκBα.

## DISCUSSION

Most NF-κB activation pathways depend on IκB kinases (Hinz & Scheidereit, 2014; Hoffmann & Baltimore, 2006). Here we provide a physiologically relevant context for IKK-independent activation of NF-κB both in human cells lines and in murine models *in vivo*. We found that in DNA damage induced senescence of epithelial cells, two interconnected events comprising a decline in IKK phosphorylation and a drop in transcription of the inhibitor of the pathway, *NFKBIA*(IκBα), initiate a persistent, IKK-independent activation of NF-κB (Fig 6).

**Figure 6.**
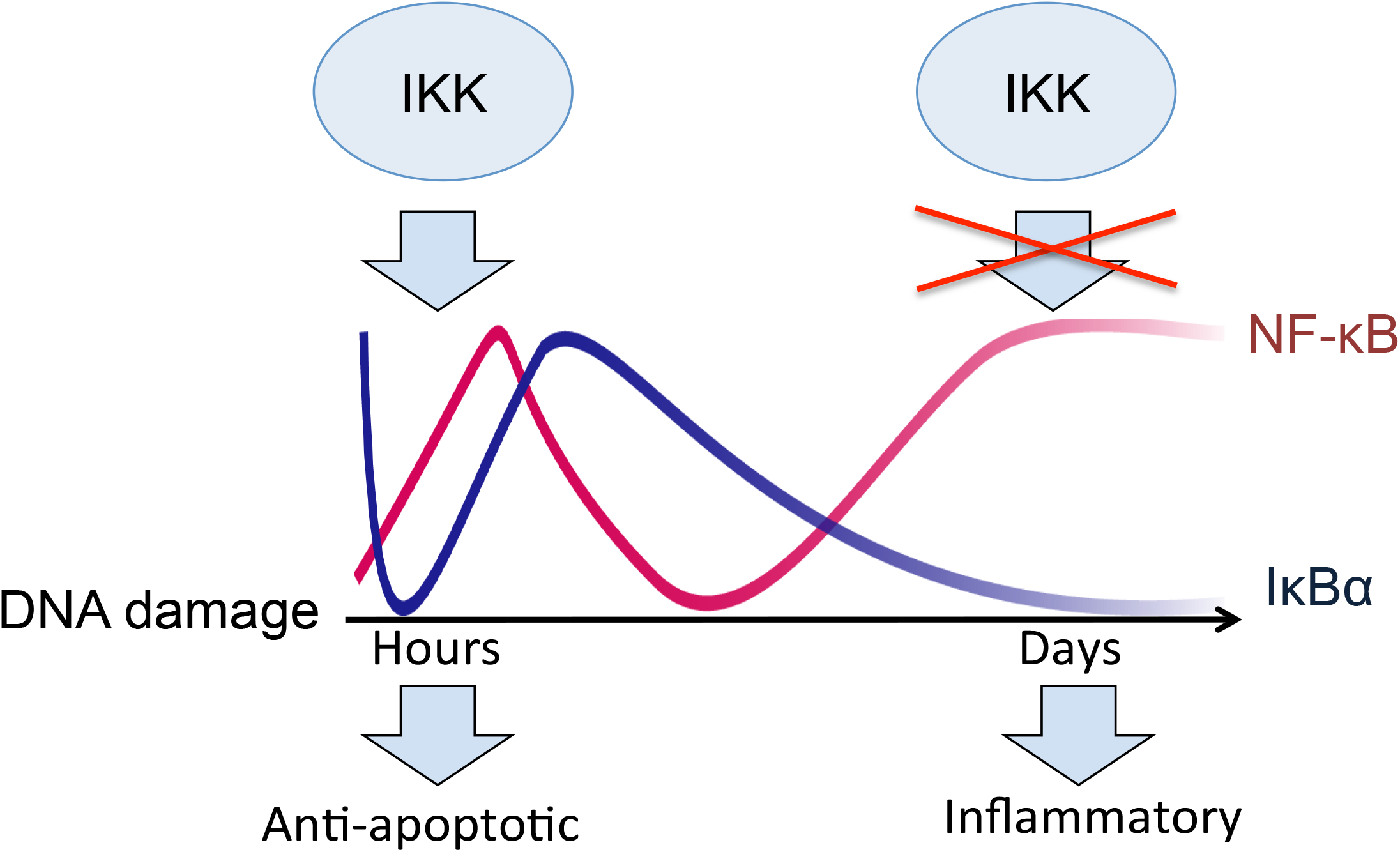
Schematic model of the two-phase NF-κB activation after DNA damage. DNA damage triggers two phases of NF-κB activation and two distinct transcriptomes: anti-apoptotic and inflammatory. NF-κB is rapidly activated through the known genotoxic stress-induced IKK cascade, resulting in proteasome-dependent IκBα degradation and expression of first phase targets genes, including IκBα. NF-κB activation in senescence is caused by a loss of IκBα expression through a largely IKK- and proteasome-independent mechanism and drives SASP expression.

We and others have previously shown that irreparable DNA damage leads to cellular senescence and to SASP driven by NF-κB (Chien et al, 2011; Jing et al, 2011; Rodier et al, 2009). However, since DNA damage triggers prompt activation of NF-κB, whose immediate transcriptional targets do not feature SASP, it was not clear how NF-κB could be responsible for two such drastically different transcriptomes arising from the same initial stimulus. Here we demonstrated both *in vivo* and *ex vivo* that a single dose of DNA damage sequentially activates two temporally and functionally distinct transcriptomes of NF-κB, separated by a span of several days. An anti-apoptotic first phase is driven by an ATM-, PARP-1- and TRAF6-dependent IKK signaling cascade (Hinz et al, 2010; Stilmann et al, 2009), resulting in proteasomal destruction of IκBα. A pro-inflammatory second phase occurs in senescence, and comprises SASP (Fig 6). Importantly, we demonstrate that the second phase of NF-κB and expression of the majority of the SASP genes are both IKK- and proteasome-independent. A fraction of transcripts that were IKK-dependent in senescence could be regulated in an alternative manner that does not require phosphorylation in the activation loop of IKKβ. Indeed, we have recently shown that basal activity of IKKβ suffices for its interaction with EDC4 (Enhancer of Decapping 4) and for post-transcriptional stabilization and destabilization of scores of transcripts, including of *CXCL8* and *TNFA (Mikuda et al, 2018)*. It is therefore possible that IKK-dependent regulation of some SASP genes occurs at the level of their RNA stability. It is also possible that additional phosphorylation sites on IKK, not analyzed in this paper, contribute to its activation and to regulation of IKK-dependent genes expressed in senescence.

Interestingly, absence of IKK at the instant of DNA damage abolishes only the first phase of NF-κB activation, but does not affect the second phase (Fig 5E). This suggests that the first phase is not required for the second, and that the changes accumulated over time activate distinct signaling pathways that enable the second phase of NF-κB. In accordance with this, we show that post-translational modifications on p65 contribute to the two distinct transcriptomes. IKK phosphorylates p65 on serine 536 within minutes following DNA damage (Fig 4), however in senescence a switch in phosphorylation from serine 536 to 468 leads to repression of *NFKBIA*. GSK3β, which is hyperactive in senescence (Ye et al, 2007), phosphorylates p65 at Ser 468. However, inhibition of GSK3β did not completely reinstate IκBα expression, indicating that additional kinases and/or epigenetic changes may contribute to downmodulation of *NFKBIA* expression in senescence.

Silencing of *NFKBIA* in senescence results in IKK-independent and persistent activation of NF-κB (Fig 6). Consequently, the second phase can be mimicked without induction of DNA damage by inactivation of the *NFKBIA* gene, as we have shown in human cell cultures and the murine knockout model (Fig 2). Nonetheless, the gene expression repertoire of *NFKBIA* knockout or knockdown cell lines does not encompass the entire senescent inflammatory response (Fig 2C and Table EV1A) likely because full-featured senescence relies on activation of additional regulators including TORC1, MAPK, Toll Like Receptors, Notch1/TGF-β, and C/EBPβ, and also on epigenetic changes that firmly establish proliferative arrest (Herranz et al, 2015; Hoare et al, 2016; Kolesnichenko et al, 2012; Narita et al, 2011; Serrano, 2012).

It was previously suggested that in addition to promoting SASP, p65 reinforces cell cycle arrest in senescent Eμ-myc lymphomas expressing the anti-apoptotic protein Bcl-2 (Chien et al, 2011). However, Bcl-2 was also shown to attenuate cell cycle progression independently of its anti-apoptotic functions (Zinkel et al, 2006). Our findings clearly demonstrate that although NF-κB negatively regulates expression of several cell cycle genes, knockdown of p65 does not disturb the established cell cycle arrest in epithelial cells and murine model described here (Table EV1A-C and data not shown). Furthermore, we did not observe establishment of senescence either in human cell lines bearing knockdown or knockout of *NFKBIA*, or in the murine mouse model. We therefore conclude that NF-κB mediates only the inflammatory phenotype, but not the cell cycle arrest in senescence.

NF-κB is constitutively activated in a variety of cancer types (Ben-Neriah & Karin, 2011). Deletion of the *NFKBIA* gene or low expression of IκBα protein serves as a major mechanism of permanent NF-κB activation in a non-classical form of glioblastoma, where it is associated with poor prognosis in patients and resistance to bortezomib and IKK inhibitors in clinical trials (Idbaih et al, 2011; Kinker et al, 2016; Raizer et al, 2016; Takeuchi & Nawashiro, 2011). Importantly, classical SASP factors and NF-κB targets, such as IL-8, IL-6 and metalloproteinases are constitutively expressed in these tumors, where they are suggested to fuel tumor growth and invasion (Puliyappadamba et al, 2014). Our results imply that in the milieu of established senescence and concomitant loss of IκBα, inhibition of the IKK-signalosome or the proteasome would be ineffective in suppressing SASP in tumor therapies.

## MATERIALS AND METHODS

### Transfection/Transduction

pTRIPz clones IκBα, p65, IKKβ, and IKKy and Scrambled or pGIPZ-IκBα and pGIPZ-scrambled (Dharmacon, Lafayette, USA) were transfected into HEK293T cells and supernatant used for transduction as described in manufacturer’s protocol (http://dharmacon.gelifesciences.com/uploadedfiles/resources/ptripz-inducible-lentiviral-manual.pdf). Clonal selection was performed using puromycin. Doxycycline hydrochloride was added daily (2 μg/ml, Sigma). Unless specified otherwise, dox treatment was done for 5-6 days prior to harvest. As controls, cells inducibly expressing scrambled shRNAs were treated with dox.

CRISPR knockout cells were generated as described previously(Mikuda et al, 2018).

*Preparation of murine tissues from in vivo experiments:* All mouse protocols in this study followed the regulatory standards of the governmental review board (Landesamt Berlin), Reg. G007/08, G 0082/13, G0358/13 and X9013/11). B6;129P2-Nfkbia^tm1Kbp^ and Tg(Vil-cre)20Syr) mice were sacrificed at 5 or 7 weeks of age. Additionally, 6 to 8 weeks old C57Bl6/N mice were sacrificed either one hour or 7 days post whole-body IR (5Gy). Control group was left untreated.

*Nuclear Cytoplasmic Fractionation, Western Blot analysis, and EMSA:* performed as described previously(Mikuda et al, 2018).

*In situ* hybridization: *was performed as described previously* (Schmidt-Ullrich et al, 2001).

*Antibody Array:* Proteome profiler (R&D Systems) antibody array was performed on 1ml of culture medium according to manufacturer’s protocol. Quantitation was performed with FusionCapt Advanced software.

*Quantitative RT-PCR* was performed using a minimum of two reference genes (TBP, Rpl13a, HRTP1) as controls, according to the manufacturer’s protocol (Promega).

*RNA-Seq:* RNA samples were prepared in quintuplicates and extracted using Trizol reagent according to manufacturer’s instructions (Thermo Fisher). Stranded mRNA sequencing libraries were prepared with 500 ng total RNA according to manufacturer’s protocol (Illumina). The libraries were sequenced in 1 x 100 +7 manner on HiSeq 2000 platform (Illumina).

## ACKNOWLEDGEMENTS

We would like to thank Wei Chen and the MDC sequencing facility for RNA-Seq analysis and Manasa Reddy Gummi, Alexandra Schulze, Lisa Spatt, Nadine Burbach and Kerstin Krueger for excellent assistance.

## REFERENCES

Acosta JC, Banito A, Wuestefeld T, Georgilis A, Janich P, Morton JP, Athineos D, Kang TW, Lasitschka F, Andrulis M, Pascual G, Morris KJ, Khan S, Jin H, Dharmalingam G, Snijders AP, Carroll T, Capper D, Pritchard C, Inman GJ, Longerich T, Sansom OJ, Benitah SA, Zender L, Gil J (2013) A complex secretory program orchestrated by the inflammasome controls paracrine senescence. Nature cell biology 15: 978–990

Acosta JC, O’Loghlen A, Banito A, Guijarro MV, Augert A, Raguz S, Fumagalli M, Da Costa M, Brown C, Popov N, Takatsu Y, Melamed J, d’Adda di Fagagna F, Bernard D, Hernando E, Gil J (2008) Chemokine signaling via the CXCR2 receptor reinforces senescence. Cell 133: 1006–1018

Bachelerie F, Ben-Baruch A, Burkhardt AM, Combadiere C, Farber JM, Graham GJ, Horuk R, Sparre-Ulrich AH, Locati M, Luster AD, Mantovani A, Matsushima K, Murphy PM, Nibbs R, Nomiyama H, Power CA, Proudfoot AE, Rosenkilde MM, Rot A, Sozzani S, Thelen M, Yoshie O, Zlotnik A (2014) International Union of Basic and Clinical Pharmacology. [corrected]. LXXXIX. Update on the extended family of chemokine receptors and introducing a new nomenclature for atypical chemokine receptors. Pharmacological reviews 66: 1–79

Beg AA, Sha WC, Bronson RT, Baltimore D (1995) Constitutive NF-kappa B activation, enhanced granulopoiesis, and neonatal lethality in I kappa B alpha-deficient mice. Genes & development 9: 2736–2746

Ben-Neriah Y, Karin M (2011) Inflammation meets cancer, with NF-kappaB as the matchmaker. Nature immunology 12: 715–723

Blagosklonny MV (2014) Geroconversion: irreversible step to cellular senescence. Cell cycle (Georgetown, Tex) 13: 3628–3635

Braig M, Lee S, Loddenkemper C, Rudolph C, Peters AH, Schlegelberger B, Stein H, Dorken B, Jenuwein T, Schmitt CA (2005) Oncogene-induced senescence as an initial barrier in lymphoma development. Nature 436: 660–665

Buss H DA, Schmitz ML, Frank R, Livingstone M, Resch K, Kracht M (2004) Phosphorylation of Serin 468 by GSK-3beta negatively regulates basal p65 NF-kappa B activity Journal of Biological Chemistry 279: 49571–49574

Chien Y, Scuoppo C, Wang X, Fang X, Balgley B, Bolden JE, Premsrirut P, Luo W, Chicas A, Lee CS, Kogan SC, Lowe SW (2011) Control of the senescence-associated secretory phenotype by NF-kappaB promotes senescence and enhances chemosensitivity. Genes & development 25: 2125–2136

Christian F SE, Carmody RJ (2016) the regulation of NF-kappa B subunits by phosphorylation Cells 5: 12

Collado M, Gil J, Efeyan A, Guerra C, Schuhmacher AJ, Barradas M, Benguria A, Zaballos A, Flores JM, Barbacid M, Beach D, Serrano M (2005) Tumour biology: senescence in premalignant tumours. Nature 436: 642

Coppe JP, Patil CK, Rodier F, Sun Y, Munoz DP, Goldstein J, Nelson PS, Desprez PY, Campisi J (2008) Senescence-associated secretory phenotypes reveal cell-nonautonomous functions of oncogenic RAS and the p53 tumor suppressor. PLoS biology 6: 2853–2868

Freund A, Patil CK, Campisi J (2011) p38MAPK is a novel DNA damage response-independent regulator of the senescence-associated secretory phenotype. EMBO J 30: 1536–1548

Hayden MS, Ghosh S (2012) NF-kappaB, the first quarter-century: remarkable progress and outstanding questions. Genes & development 26: 203–234

Herranz N, Gallage S, Mellone M, Wuestefeld T, Klotz S, Hanley CJ, Raguz S, Acosta JC, Innes AJ, Banito A, Georgilis A, Montoya A, Wolter K, Dharmalingam G, Faull P, Carroll T, Martinez-Barbera JP, Cutillas P, Reisinger F, Heikenwalder M, Miller RA, Withers D, Zender L, Thomas GJ, Gil J (2015) mTOR regulates MAPKAPK2 translation to control the senescence-associated secretory phenotype. Nature cell biology 17: 1205–1217

Hinz M, Arslan SC, Scheidereit C (2012) It takes two to tango: IkappaBs, the multifunctional partners of NF-kappaB. Immunological reviews 246: 59–76

Hinz M, Scheidereit C (2014) The IkappaB kinase complex in NF-kappaB regulation and beyond. EMBO reports 15: 46–61

Hinz M, Stilmann M, Arslan SC, Khanna KK, Dittmar G, Scheidereit C (2010) A cytoplasmic ATM-TRAF6-cIAP1 module links nuclear DNA damage signaling to ubiquitin-mediated NF-kappaB activation. Molecular cell 40: 63–74

Hoare M, Ito Y, Kang TW, Weekes MP, Matheson NJ, Patten DA, Shetty S, Parry AJ, Menon S, Salama R, Antrobus R, Tomimatsu K, Howat W, Lehner PJ, Zender L, Narita M (2016) NOTCH1 mediates a switch between two distinct secretomes during senescence. Nature cell biology 18: 979–992

Hoffmann A, Baltimore D (2006) Circuitry of nuclear factor kappaB signaling. Immunological reviews 210: 171–186

Idbaih A, Marie Y, Sanson M (2011) NFKBIA deletion in glioblastomas. The New England journal of medicine 365: 277; author reply 277-278

Jing H, Kase J, Dorr JR, Milanovic M, Lenze D, Grau M, Beuster G, Ji S, Reimann M, Lenz P, Hummel M, Dorken B, Lenz G, Scheidereit C, Schmitt CA, Lee S (2011) Opposing roles of NF-kappaB in anti-cancer treatment outcome unveiled by cross-species investigations. Genes & development 25: 2137–2146

Kinker GS, Thomas AM, Carvalho VJ, Lima FP, Fujita A (2016) Deletion and low expression of NFKBIA are associated with poor prognosis in lower-grade glioma patients. Scientific reports 6: 24160

Klement JF, Rice NR, Car BD, Abbondanzo SJ, Powers GD, Bhatt PH, Chen CH, Rosen CA, Stewart CL (1996) IkappaBalpha deficiency results in a sustained NF-kappaB response and severe widespread dermatitis in mice. Molecular and cellular biology 16: 2341–2349

Kolesnichenko M, Hong L, Liao R, Vogt PK, Sun P (2012) Attenuation of TORC1 signaling delays replicative and oncogenic RAS-induced senescence. Cell cycle (Georgetown, Tex) 11: 2391–2401

Krappmann D, Wulczyn FG, Scheidereit C (1996) Different mechanisms control signal-induced degradation and basal turnover of the NF-kappaB inhibitor IkappaB alpha in vivo. The EMBO journal 15: 6716–6726

Kuilman T, Michaloglou C, Vredeveld LC, Douma S, van Doorn R, Desmet CJ, Aarden LA, Mooi WJ, Peeper DS (2008) Oncogene-induced senescence relayed by an interleukin-dependent inflammatory network. Cell 133: 1019–1031

Lasry A, Ben-Neriah Y (2015) Senescence-associated inflammatory responses: aging and cancer perspectives. Trends Immunol 36: 217–228

Lee S, Schmitt CA (2019) The dynamic nature of senescence in cancer. Nature cell biology 21: 94–101

Mikuda N, Kolesnichenko M, Beaudette P, Popp O, Uyar B, Sun W, Tudan AB, Perder B, Akalin A, Chen W, Mertins P, Dittmar G, Hinz M, Scheidereit C (2018) The IkappaB kinase complex is a regulator of mRNA stability. The EMBO journal 14: 24

Milanovic M, Fan DNY, Belenki D, Dabritz JHM, Zhao Z, Yu Y, Dorr JR, Dimitrova L, Lenze D, Monteiro Barbosa IA, Mendoza-Parra MA, Kanashova T, Metzner M, Pardon K, Reimann M, Trumpp A, Dorken B, Zuber J, Gronemeyer H, Hummel M, Dittmar G, Lee S, Schmitt CA (2018) Senescence-associated reprogramming promotes cancer stemness. Nature 553: 96–100

Narita M, Young AR, Arakawa S, Samarajiwa SA, Nakashima T, Yoshida S, Hong S, Berry LS, Reichelt S, Ferreira M, Tavare S, Inoki K, Shimizu S, Narita M (2011) Spatial coupling of mTOR and autophagy augments secretory phenotypes. Science (New York, NY) 332: 966–970

Puliyappadamba VT, Hatanpaa KJ, Chakraborty S, Habib AA (2014) The role of NF-kappaB in the pathogenesis of glioma. Molecular & cellular oncology 1: e963478

Raizer JJ, Chandler JP, Ferrarese R, Grimm SA, Levy RM, Muro K, Rosenow J, Helenowski I, Rademaker A, Paton M, Bredel M (2016) A phase II trial evaluating the effects and intra-tumoral penetration of bortezomib in patients with recurrent malignant gliomas. Journal of neuro-oncology 129: 139–146

Reimann M, Lee S, Loddenkemper C, Dorr JR, Tabor V, Aichele P, Stein H, Dorken B, Jenuwein T, Schmitt CA (2010) Tumor stroma-derived TGF-beta limits myc-driven lymphomagenesis via Suv39h1-dependent senescence. Cancer cell 17: 262–272

Rodier F, Coppe JP, Patil CK, Hoeijmakers WA, Munoz DP, Raza SR, Freund A, Campeau E, Davalos AR, Campisi J (2009) Persistent DNA damage signalling triggers senescence-associated inflammatory cytokine secretion. Nature cell biology 11: 973–979

Salama R, Sadaie M, Hoare M, Narita M (2014) Cellular senescence and its effector programs. Genes & development 28: 99–114

Schmidt-Ullrich R, Aebischer T, Hulsken J, Birchmeier W, Klemm U, Scheidereit C (2001) Requirement of NF-kappaB/Rel for the development of hair follicles and other epidermal appendices. Development (Cambridge, England) 128: 3843–3853

Serrano M (2012) Dissecting the role of mTOR complexes in cellular senescence. Cell cycle (Georgetown, Tex) 11: 2231–2232

Shelton DN, Chang E, Whittier PS, Choi D, Funk WD (1999) Microarray analysis of replicative senescence. Current biology : CB 9: 939–945

Smale ST (2012) Dimer-specific regulatory mechanisms within the NF-kappaB family of transcription factors. Immunological reviews 246: 193–204

Stilmann M, Hinz M, Arslan SC, Zimmer A, Schreiber V, Scheidereit C (2009) A nuclear poly(ADP-ribose)-dependent signalosome confers DNA damage-induced IkappaB kinase activation. Molecular cell 36: 365–378

Takeuchi S, Nawashiro H (2011) NFKBIA deletion in glioblastomas. The New England journal of medicine 365: 276–277; author reply 277–278

Wu ZH, Shi Y, Tibbetts RS, Miyamoto S (2006) Molecular linkage between the kinase ATM and NF-kappaB signaling in response to genotoxic stimuli. Science (New York, NY) 311: 1141–1146

Ye X, Zerlanko B, Kennedy A, Banumathy G, Zhang R, Adams PD (2007) Downregulation of Wnt signaling is a trigger for formation of facultative heterochromatin and onset of cell senescence in primary human cells. Molecular cell 27: 183–196

Zinkel S, Gross A, Yang E (2006) BCL2 family in DNA damage and cell cycle control. Cell death and differentiation 13: 1351–1359

